# Are sleep spindles poised on supercritical Hopf bifurcations?

**DOI:** 10.1101/512145

**Authors:** M. Ospeck

**Affiliations:** Mathematics Department, Suny Poly, Utica, NY, US

## Abstract

Sleep spindles are recognized as an important intermediate state of long term memory formation. During non REM sleep, large numbers of thalamic relay neurons synchronize their spike bursts for one half to two seconds, entraining many millions of neurons, and constituting a sleep spindle. Here we study spindle amplification, entrainment, synchronization and decay. Relay neurons have both a high resting state near −60 millivolts (mV) and low resting state near −75 mV. Due to the neuron’s sodium conductance, low-threshold calcium conductance, and calcium-dependent H conductance, it exhibits a number of bifurcations, like its supercritical Hopf at −61 mV. Here low-threshold calcium conductance destabilizes membrane potential to birth a small limit-cycle in the 7-16 Hz range. Supercritical Hopf bifurcations are the underlying mechanism for amplification and frequency selectivity in hearing: hair cells are forced by sinusoidal input currents driving their mainly capacitive loads, with the forcing currents locking at 90 degree phase leads with respect to their oscillating membrane potentials. Here we model a small part of a spindle, with 6 cross-coupled relay neurons all poised on Hopf bifurcations. One neuron is forced by a weak noisy train of periodic current impulses that typically lock at a 90 degree phase lead with respect to its voltage oscillation. It then drives its neighbors, causing them to drive each other at much smaller phase angles, usually less than ±10 degrees. The system of Hopf oscillators exhibit small signal amplification and frequency selectivity, high degrees of synchronization and noise rejection, and switch-ability. These argue in favor of spindling relay neurons poising on, or very near to, supercritical Hopf bifurcations. Also, during the phase-locking of their spike bursts, calcium conductance oscillations increase internal calcium, which turns on slow H current. This depolarizes the relay cells, pushing them below their Hopfbifurcations and terminating the spindle.

## INTRODUCTION

Mainly all conscious perception passes through the thalamus on its way to the cortex [1]. Thalamic relay neurons, acting in their relay mode, forward rate-coded spike trains from subcortical sensory areas through specific thalamic relay nuclei to their related areas of cortex. Examples are cochlea → medial geniculate body (MGB) → auditory cortex, and retina → lateral geniculate nucleus (LGN) → visual cortex, where the arrows stand for tonic firing sensory neurons and tonic firing relay neurons, respectively.

Relay neurons have both strong driver and weaker modulator inputs [1]. Small numbers of drivers from, for example, retinal ganglion cells, or layer V cortex pyramid neurons, make large glomerulus-type synapses onto the proximal dendrites and somas of thalamic relays [1]. On the other hand, large numbers of modulator inputs from layer VI cortex pyramid neurons, thalamic interneurons, or parabrachial neurons, make weaker synaptic contacts onto relay neuron distal dendrites [1]. It should be noted that thalamic reticular nucleus neurons differ in that they are able to make both strong and weak inputs onto relay neurons [1]. In the case of vision, only 7% of the LGN relay inputs are retinal drivers, but these define the receptive field properties of the visual cortex target [1]. This is an example of a general principle called “labeled line,” in which a sparse number of strong sensory driver inputs are able to control the receptive field properties of the cortical target area [1].

In addition to the driver vs. modulator input distinction, relay neurons can be partitioned into first order vs. higher order [1]. Retina → LGN → visual cortex involves first order (FO) relay neurons that advance awake state visual information. Contrast this against the higher order (HO) relays, located mainly in the thalamic pulvinar nucleus. Instead of forwarding sensory information, HO relays pass spike trains from a cortex source to another cortex target [1]. The pulvinar is the largest region of the thalamus and most of the thalamus is devoted to HO relays [1]. For example, a layer V pyramid in cortex region 1 drives an HO pulvinar relay that in turn drives cortex region 2, whose layer V pyramid then drives another HO pulvinar relay making inputs to cortex region 3, etc. [1]. Such cortex → pulvinar → cortex → pulvinar →…HO relay pathways are important in the awake state and also during non-rapid eye movement (NREM) sleep for the propagation of sleep spindles, delta waves and slow waves [1].

Besides their tonic-firing relay mode, thalamic relay cells have a burst firing mode that is used for both attention and memory. For example take walking late at night along a dark country road. One might be startled to see a flaming meteor whiz past, only to realize that it was just a firefly flying close by your right ear. Burst firings by hyperpolarized thalamic relays are part of this attention-grabbing effect. However, relay neurons have a number of distinct burst modes, several of which are used during NREM sleep for memory reprocessing [2]. NREM slow wave sleep includes sleep spindles (7-16 Hz), delta waves (δ; 1-4 Hz), and slow waves (SW; 0.3-1 Hz) [2]. Interestingly, it also contains brief episodes of high-frequency beta (β; 13-30 Hz) and gamma (γ; 30-60 Hz) that are normally associated with the awake state [2].

Repeated episodes of synchronized slow wave activity are thought to lock in declarative memories [2]. During the awake state, neocortex plays a version of the day’s events to the hippocampus. Then during slow wave sleep, the hippocampus recapitulates these short term memories back to the cortex. This playback involves short intervals of high frequency β and γ, and it is required to construct a permanent memory in cortex, which subsequently becomes independent of the hippocampus [2]. Destexhe and Sejnowski’s “recall-store” memory consolidation hypothesis is based on brief episodes of 7-16 Hz spindles and β and γ from the hippocampus simultaneously driving cortex targets, with these events followed by slow waves [2]. During this part of slow wave sleep the δ and SW force the same cortical targets, but in a lower < 4 Hz frequency range [1, 2, 3, 4]. They conclude that slow wave sleep appears to be a cyclic, iterative process that leads to “off line” memory reprocessing by driving cortex pyramid neurons in complementary ways [2]. Additionally, more than just reactivating memory traces, slow wave sleep increases signal-to-noise-ratio on Py dendrites by strengthening some synapses and downregulating the weights of others, effectively turning some of them from the active to the latent state [2, 5]. Everyone has had the experience of sleeping on a problem, which then becomes clear in the morning.

Cortex pyramid neurons (Py) have a hyperpolarized down-state able to cause a local cortex down-state (low β, γ, but high δ, SW) [4]. Coincident down-states from different cortex regions are then able to initiate a focal thalamic down-state that births a spindle [3, 4]. The nomenclature for up-states and down-states can be confusing: another way of saying the same thing is that the on-set of a cortex up-state sends a burst of spikes to reticular nucleus neurons, which then hyperpolarize relays, causing them to initiate a spindle [5]. Subsequently these regions receive the feedback tetanus from the spindle together with a high-frequency playback of a part of the day’s short-term memories from the hippocampus, all this typically occurring during their down-state to up-state transitions [4]. Surprisingly, during these episodes some cortex Py neuron targets are still able to maintain relatively low firing rates because the spindle simultaneously forces EPSPs (excitatory post synaptic potentials) in their dendrites, and larger IPSPs (inhibitory post synaptic potentials) in their somas [2]. Py dendrites are known to be dominated by excitatory synapses, while their somas have mainly inhibitory ones [2]. In this way 7-16 Hz spindles are able to activate NMDARs (N-methyl-D-aspartate receptors) that bring calcium into Py excitatory synapses, while at the same time acting as a governor on the Py spike rate [2]. It’s also thought that the spindle frequency range is optimal for the activation of calcium-modulated kinase 2 (CaMKII), which is known to effect long term potentiation (LTP) at excitatory synapses [2]. Also activated is protein kinase A (PKA) which is known to inhibit nuclear phosphatases that block gene expression [2]. “Recall-store” considers synaptic calcium infused by a spindle as being able to “tag” a particular Py synapse for subsequent long-term changes [2]. Then after the spindle, calcium induced calcium release (CICR) from endoplasmic reticulum into the nucleus during δ and SW would drive a calcium-dependent gene expression, sensitive to slow waves, that is able to transcribe channel proteins destined for a tagged synapse [2]. It is also known that subthreshold inputs to Py neurons during their up-state induces synaptic weakening, but that weakening can be prevented by correlated postsynaptic spiking, with synaptic protection dependent on NMDA channels and glycogen synthase kinase (GSK3β) [6]. It appears that slow wave sleep defaults towards synaptic depression and that active defense is required to maintain or increase synaptic weights.

Spindle-like oscillations have previously been modelled based on poising relays on subcritical Hopfbifurcations [7]. A second model involves a relay cell strongly coupled to a reticular nucleus neuron as being the effective oscillatory unit [8]. Spindles occur in neocortex, the parahippocampal gyrus and the hippocampus, and originate thalamically, while down-states originate cortically, with thousands of thalamic spindles and cortex down-states occurring during a typical night of sleep [3, 4]. There is a sharp transition around the supplementary motor area between fast centroparietal spindles (13-15 Hz), often occurring with cortex slow wave up-states, and slower frontal spindles (9-12Hz) that occur 200 ms later on average [3]. Normally spindles start at the thalamic down-state peak and end near its following upstate peak [4]. A spindle waxes as more relay neurons become entrained and phase-lock their spikes (amplification phase), then plateaus (with large numbers of neurons being entrained), and subsequently wanes as phase-locking is lost (decay phase). Here we consider a higher frequency (15 Hz) spindle in order to study spindle creation, amplification, entrainment and decay. We use a simple model of small group of coupled relay neurons to investigate the advantages for poising these on, or very near to, supercritical Hopfbifurcations.

## MODEL

The sleep spindle model used in this study employs the Wang model of a thalamic relay neuron [9] and includes the calcium-dependence of its H current [7, 10]. Wang’s membrane potential charging equation (Eq.1) has 7 currents: 3 high voltage-threshold ones: the inactivating sodium current *I_Na_*, the persistent sodium current *I*_*Na*(*p*)_, and the delayed rectifier potassium current *I_k_*. Together with linear leak current *I_L_* these make a Hodgkin-Huxley action potential generator that spikes spontaneously when depolarized above −56 mV. Two low voltage activated currents, the low-threshold calcium current *I_T_* and the hyperpolarization-activated “sag” current *I_H_*, along with *I_L_*, comprise a low frequency (16.5 Hz) subthreshold oscillator that is active below −60 mV. Externally-applied bias current *I_app_* stands in for modulator-type synaptic inputs [1] from layer VI cortex neurons, thalamic reticular nucleus neurons, parabrachial neurons, and thalamic interneurons, all of which are able to move the relay neuron’s voltage operating point *v*.

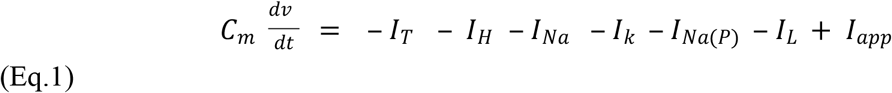

The model has a number of interesting bifurcations and rest states that are functions of membrane potential [9]. There is a saddle-node bifurcation near −55 mV, a high voltage resting state centered at −60.5 mV, a supercritical Hopf bifurcation that births a small 16.5 Hz limit cycle, near −63 mV, and a fold-circle bursting bifurcation several tenths of a mV below that. The bursts become stronger, but lower in frequency, until a low voltage resting state is achieved near −76 mV.

With 3 exceptions, we use Wang’s original parameters. In order to be consistent with the frequency range associated with sleep spindles (7-16 Hz) [2], (9-15 Hz) [3] we lengthened the time constant on the *h* off-gate of the low-threshold calcium conductance. The super critical Hopf bifurcation now occurs at −61 mV, where it births a small 15 Hz limit-cycle. Also, as per [7] we made the two H conductance activation gates *H_s_* and *H_f_* both calcium-sensitive [10]. Wang’s *H* gates had a similar voltage-dependent activation and slow (~1 sec) voltage-dependent time constant as [7] faster *H_f_* gate. The very slow *H_s_* gate was included based on [7] observation that in order to explain the H channel on→ off and off→ on transitions one of the gates had to be significantly slower in the higher voltage range. Membrane capacitance *C_m_* is 1.0 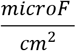 currents are in 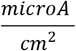, conductances in 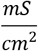, and voltages in mV. Eq.2 shows relay neuron N1’s charging currents, conductances, gating variables and voltage drives. It includes synaptic inputs *I_s_* from N1’s neighbors and externally:

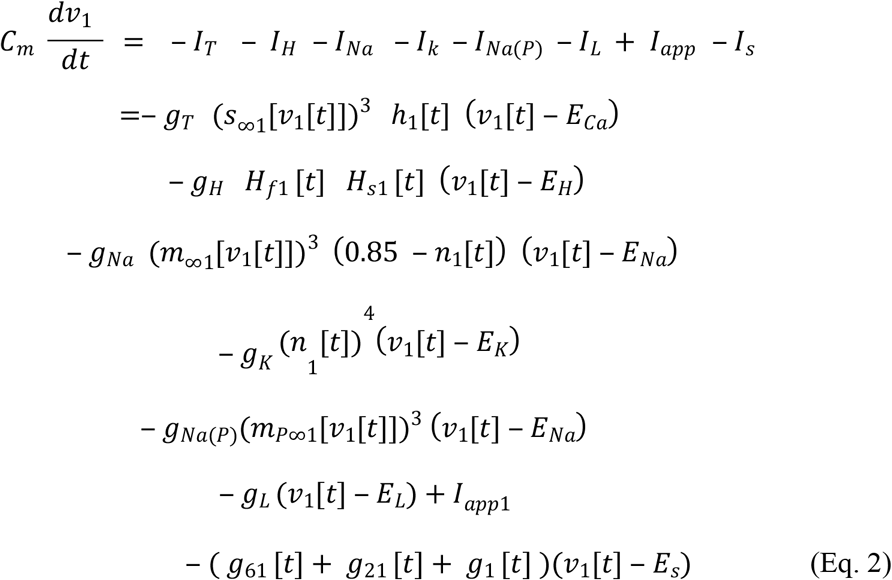

N1 has 6 dynamical variables: *v*_1_[*t*], its membrane potential, *h*_1_[*t*], the relatively slow voltage-dependent off gate on its low threshold calcium conductance, *H*_*f*1_ [*t*] and *H*_*s*1_ [*t*], the faster and slower voltage-, and calcium-, dependent activation gates of its very slow hyperpolarization-activated “sag” current, *n*_1_[*t*] the voltage-dependent activation gate of the delayed rectifier potassium conductance, and *Ca*_1_[*t*], the calcium concentration near the inner membrane where the calcium-sensitive H conductance is located. Very fast gates such as *s*, the activation gate of the calcium conductance, *m*, the activation gate of the inactivating sodium conductance, and *m_P_* the activation gate of the persistent sodium conductance, are treated as instantaneous functions of membrane potential [9]. The off gate on the inactivating sodium conductance was replaced by (0.85 - *n*_1_[*t*]) according to Fitzhugh’s observation of it having a similar time course and linear relationship with the potassium conductance activation gate [9]. *g*_61_ [*t*] represents synaptic input conductance from neuron N6 to N1, *g*_21_ [*t*] from N2 to N1. *g*_1_ [*t*] is from a weak, noisy periodic template that is used to mimic external synaptic input from a layer V pyramid neuron.

*I_app_* is used as a control parameter to change the voltage operating point [9]. Lacking synaptic input, *I_app_* depolarizations that displace membrane potential above −55 mV result in an increasing rate of spontaneous spiking. This is typical behavior for a saddle-node on invariant circle (SNIC) bifurcation [11, 12]. Hyperpolarizing currents that pull membrane potential down from the −60 mV resting state to −61 mV generate a small 15 Hz sinusoidal oscillation (supercritical Hopf bifurcation). Additional hyperpolarization increases the size of this oscillation, until spontaneous spiking starts at around −61.3 mV (fold-circle bursting bifurcation) [12]. Continuing to hyperpolarize below the bursting bifurcation increases the number of sodium spikes in the bursts that ride the peaks of the calcium-driven limit cycle [9]. Hyperpolarization also slows down the frequency of the bursts from 15 Hz to 1 Hz, prior to them disappearing entirely at −75 mV, the low voltage resting state. The voltage range of interest for this study is near to the supercritical Hopf bifurcation, −60 to −61.2 mV. In this region the input conductance 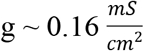 so that a change in *I_app_* by 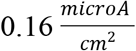 displaces the voltage operating point by ~1 mV (*I* = *g v*), and the effective membrane time constant 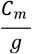 is ~ 6 ms.

The Mathematica model in the supplementary file lists parameter values and has 6 cross-coupled relay neurons with external forcing of N1. It’s based on the following assumptions:

1. It assumes a higher order (HO) thalamic pulvinar relay from one cortex region drives a second cortex patch whose layer V pyramid then drives another HO pulvinar relay cell, etc. [1]. It presupposes sequences like cortex1 layer V pyramid → pulvinar relay 2 → cortex2 layer V pyramid→ pulvinar relay 3 etc.
2. It assumes a low latency pathway from relay 1 to relay 2 due to fast myelinated nerve fiber conduction velocities 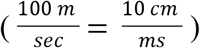 [13]. This pathway employs large, fast, glomerulus-type, driver synaptic connections between layer V pyramid neurons and relay neurons [1]. Hearing is an example of fast nerve fiber conduction and fast synapses: spike trains in primary auditory neurons in guinea pigs are able to maintain phase locking to a mechanical sound wave up to about 3 kHz (in cats, up to 6 kHz) [14]. We assume 0.5 - 1.0 ms latency between a spike in one HO relay neuron and its EPSC in a second relay. As they relate to synch index [15], latencies from 0.5 to 4.0 ms are investigated in Fig. 4.
3. Assumes cortex down-states (low β, γ, but high δ, SW in power spectrum) are via bias currents able to entrain particular relay cells [4]. Coincident cortical down-states are able to drive specific thalamic reticular nucleus neurons which then hyperpolarize the membrane potential of particular relay neurons via fast GABA A and slower, stronger GABA B synaptic inputs [1, 4]
4. For the purposes of illustrating spindle propagation and entrainment a simple ring geometry with 6 cross-coupled relay neurons was chosen. Only one of them (N1) is driven by a noisy periodic template intended to mimic external synaptic input from a layer V pyramid neuron.
5. The size of the synaptic AMPA conductance due to a single presynaptic vesicle release (mini) was chosen to be 0.1 *mS*, since this resulted in large EPSPs in the 1-2 mV range. Synaptic transmission is unreliable, caused mainly by probability release mechanisms at low capacity synapses [16]. In the weak forcing regime each N1 external presynaptic spike releases a Poisson average of 0.5 vesicles, where a single vesicle (mini) makes a ~1.5 mV post synaptic potential.
6. Assumes fast calcium clearance away from the relay neuron’s inner membrane’s calcium-sensitive H conductance via diffusion, chelators and pumps. Investigated here are clearance rates of 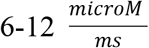. We use the hair cell as an example, since it has a very high density of calcium pumps in its hair bundle and several mM concentrations of various calcium buffers in its soma, both of these able to rapidly lower its internal calcium concentration (outer hair cells have similar high calcium chelator concentrations as do muscle fibers) [17]. Previously [7] used a very low calcium-pump-only clearance rate of 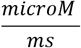.
7. Investigates the likely range from 0.5 to 10.0 *microM* for the calcium sensitivity threshold of the H conductance [7].

## RESULTS

We use 6 coupled relay neurons to investigate the consequences of them being poised on supercritical Hopf bifurcations, all with the same center frequency (CF), for the purposes of making and propagating highly synchronized sleep spindles. We force one of them, N1, with a weak noisy periodic sequence of short current impulses designed to mimic EPSCs (excitatory post synaptic currents).

Membrane potential is a natural control parameter for adjusting proximity to a supercritical Hopf bifurcation in an electrically excitable cell [11, 12]. We use bias current *I_app_* as a control parameter to move the neuron’s voltage operating point with respect to the bifurcation [9]. In addition, the cell has a calcium-sensitive H current, where its internal calcium, acting as a feedback, is able to readjust its membrane potential with respect to the bifurcation. Fig. 1 shows two sample runs, the first at high gain, and the second at low gain, both with respect to calcium influence on H current. At the start of a simulation *I_app_* is used to poise the model neurons on supercritical Hopfbifurcations with a CF of 15 Hz. It stands in for modulator-type bias currents, the most important of which is from thalamic reticular nucleus neurons that make synapses onto relay neurons able to activate GABA A and GABA B receptors [1]. GABA (gamma-Aminobutyric acid) binding to the A receptor rapidly turns on a chloride channel that hyperpolarizes the neuron [1]. B receptor activation is more powerful and long-lasting, activating a second messenger pathway that slowly turns on a number of potassium currents that more strongly hyperpolarize the neuron [1].

**Fig.1.**
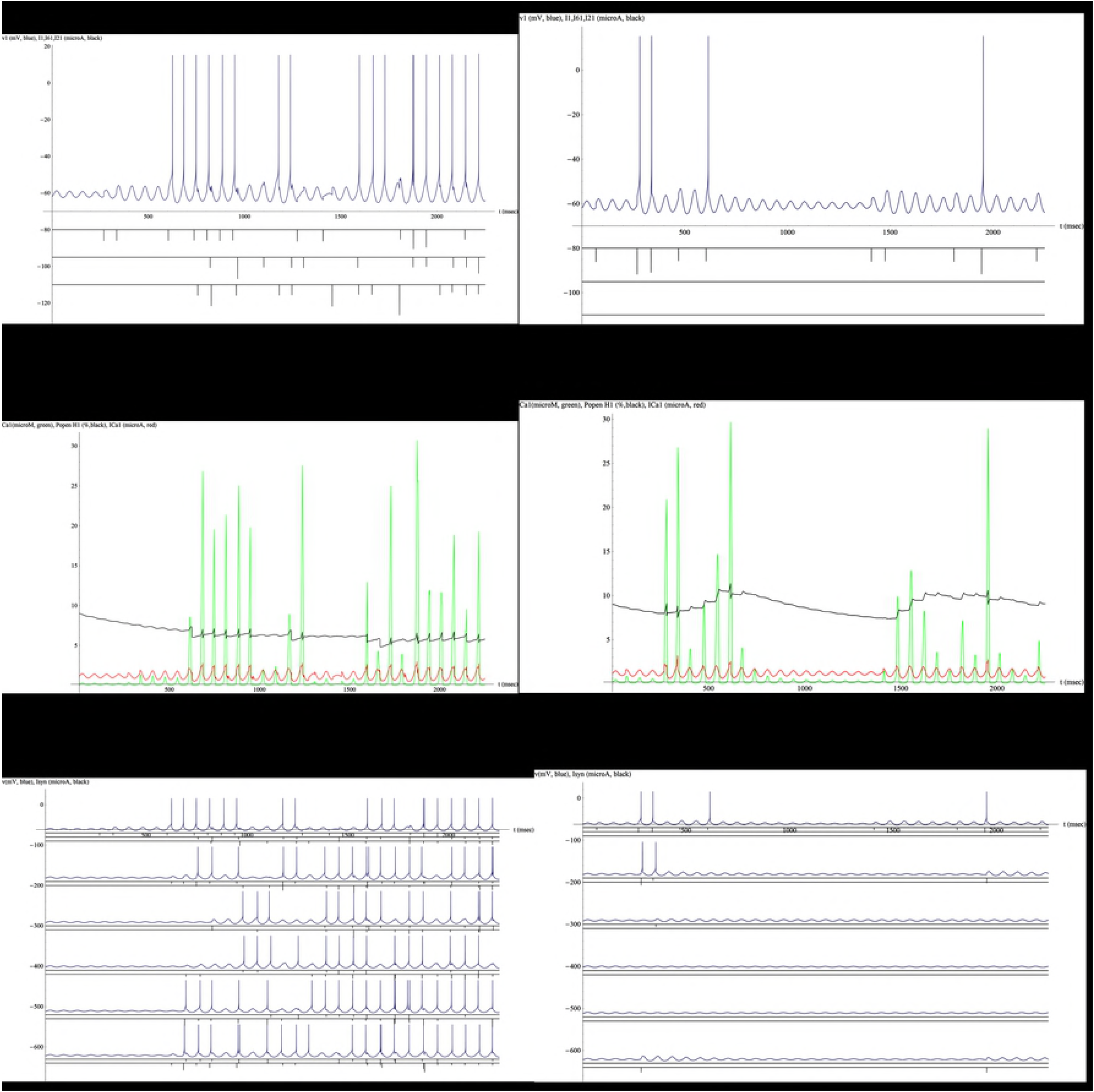
Forcing relay neurons by a weak noisy series of periodic current impulses. A. Neuron N1 is forced at its center frequency (CF=15 Hz). All 6 cross-coupled relays are poised on supercritical Hopf bifurcations with the same CF. 13 forcing EPSCs result in 18 N1 spikes. High gain is due to a high calcium clearance rate of 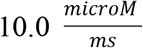 and high threshold H current calcium sensitivity of 2.0 *microM*. N1 membrane potential (blue), synaptic current impulses (black), external, N6 → N1 and N2 → N1 in descending order. B. N1 low-threshold calcium current (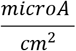, red), calcium concentration near H channels (*microM*, green) and open probability of voltage- and calcium-sensitive H current (%, black). Internal calcium turns on H, depolarizing the neurons, turning off calcium conductance, driving them below the bifurcation and lowering gain. C. 106 spikes are shown in descending order from N1 to N6 (blue), along with the synaptic EPSCs exchanged between them (black), for example N2: N1 → N2 above N3 → N2. Spike synchronization is high (average spike synch index 0.5). D. N1 spikes for same forcing, but low gain, i.e. the neurons have a lower calcium clearance rate of 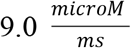and a lower H calcium sensitivity threshold of 1.0 *microM*. E. Increased H channel open probability (contrast with B). F. Low spike rate (contrast with C).

In the high gain example (Fig. 1 A, B, C) internal calcium from calcium conductance oscillations is not much able to turn on H currents and raise the relay neurons’ membrane potentials. Higher membrane potentials near −60 mV almost completely turn off calcium conductance. Hence in this case the relay neurons remain close to the Hopf bifurcation at −61 mV and respond with a high spike rate. Contrast this against D, E, F where calcium is able to turn on H current and depolarize the relay neurons away from the bifurcation, causing their spike output to plummet. In plate D, in between forcing currents (600 to 1400 ms) the decay of N1’s membrane potential oscillation is clear. This lower excitability is due to increased open probability of H conductance, causing a depolarization that inactivates calcium conductance (plate E; 600 ms).

In Fig. 2 we give an overview of spike-gain parameter space that involves calcium-clearance and the calcium-sensitivity threshold of the H current. Feedback by H depolarizes the cell below the Hopf bifurcation, lowering its excitability and spike output.

**Fig.2.**
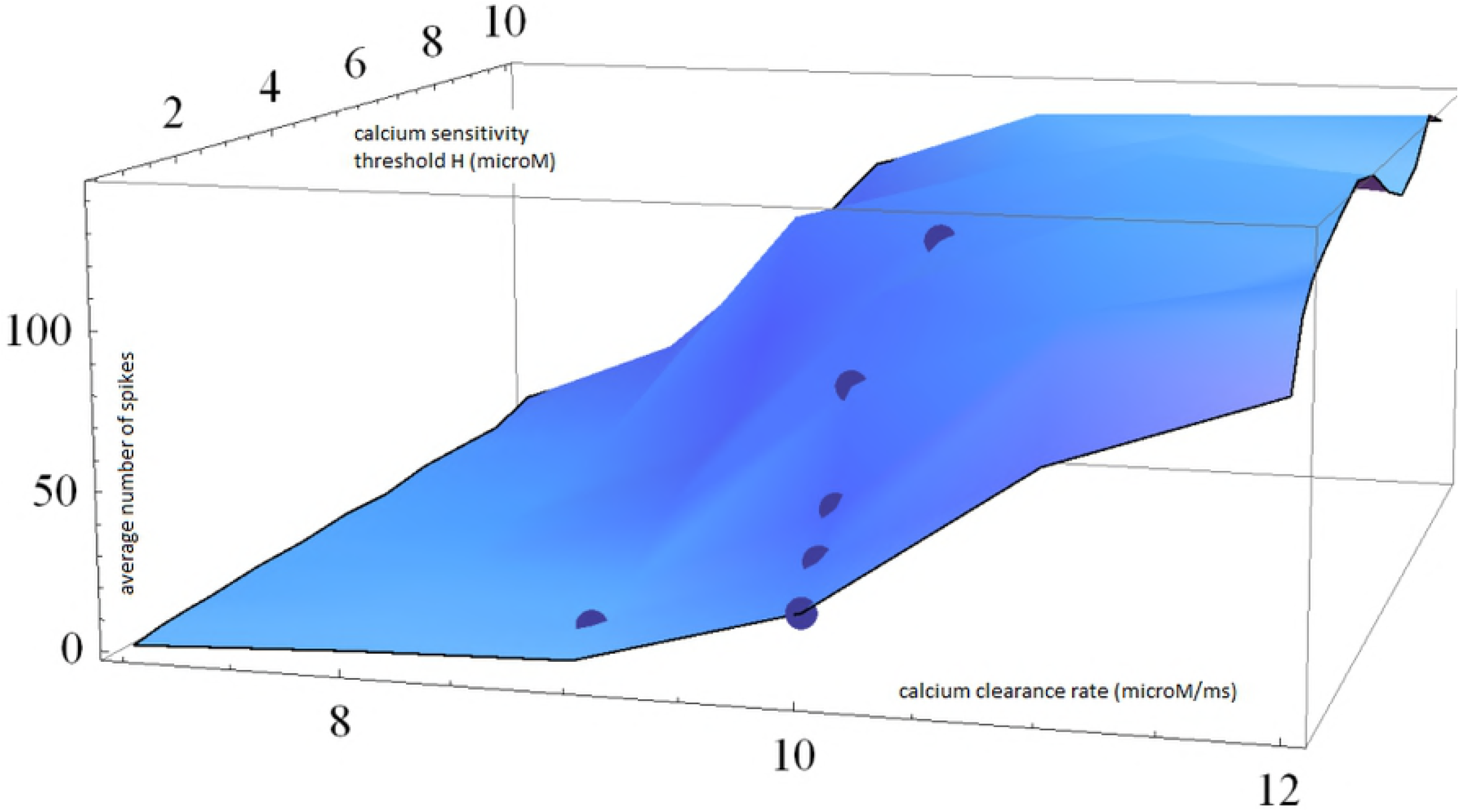
Spike output for neurons N1-N6 as a function of the calcium clearance rate near the membrane and the calcium sensitivity threshold of the H channels. More calcium or a lower sensitivity threshold more quickly depolarizes the relays, driving them below the Hopf bifurcation. Average spike output for (*CaClear, CaSensH*) = (9.0, 1.0), (10.0, 0.5), (10.0, 1.0), (10.0, 1.5), (10.0, 2.0) and (10.0, 5.0) are shown as dots.

Depth EEG (electroencephalogram), also known as SEEG (stereo EEG), positions electrodes near internal brain structures like thalamus, hippocampus or cortex, so is able to record close-in field potentials from groups of neurons [3, 4]. Fig. 3 A is a cartoon of a depth EEG recording showing the collective mode of a large number of pulvinar relays participating in a sleep spindle. The figure was drawn based on experimental recordings, and shows a local thalamic down-state inducing a spindle [3, 4]. Note that cortex down-states usually precede a thalamic down-state, and then the locally hyperpolarized thalamus (down-state) launches the spindle. We model a spindle focus in plates B, C, D where spiking is initiated by self-excitation, rather than by driver synaptic inputs, and is subsequently terminated by internal calcium. In plate B the model relays start out hyperpolarized, overexcited by modulator input to slightly above the Hopf bifurcation, near the fold-circle bursting bifurcation. But while the neurons start out in an overexcited state, they are relatively sensitive to internal calcium. What results is a case of self-excitation via bias currents, then spindle termination due to calcium activation of H current. Note that small voltage oscillations increase H open probability, while spiking lowers it. This is due to the nature of the H “sag current,” in that it is turned on by calcium and off by depolarization.

**Fig.3.**
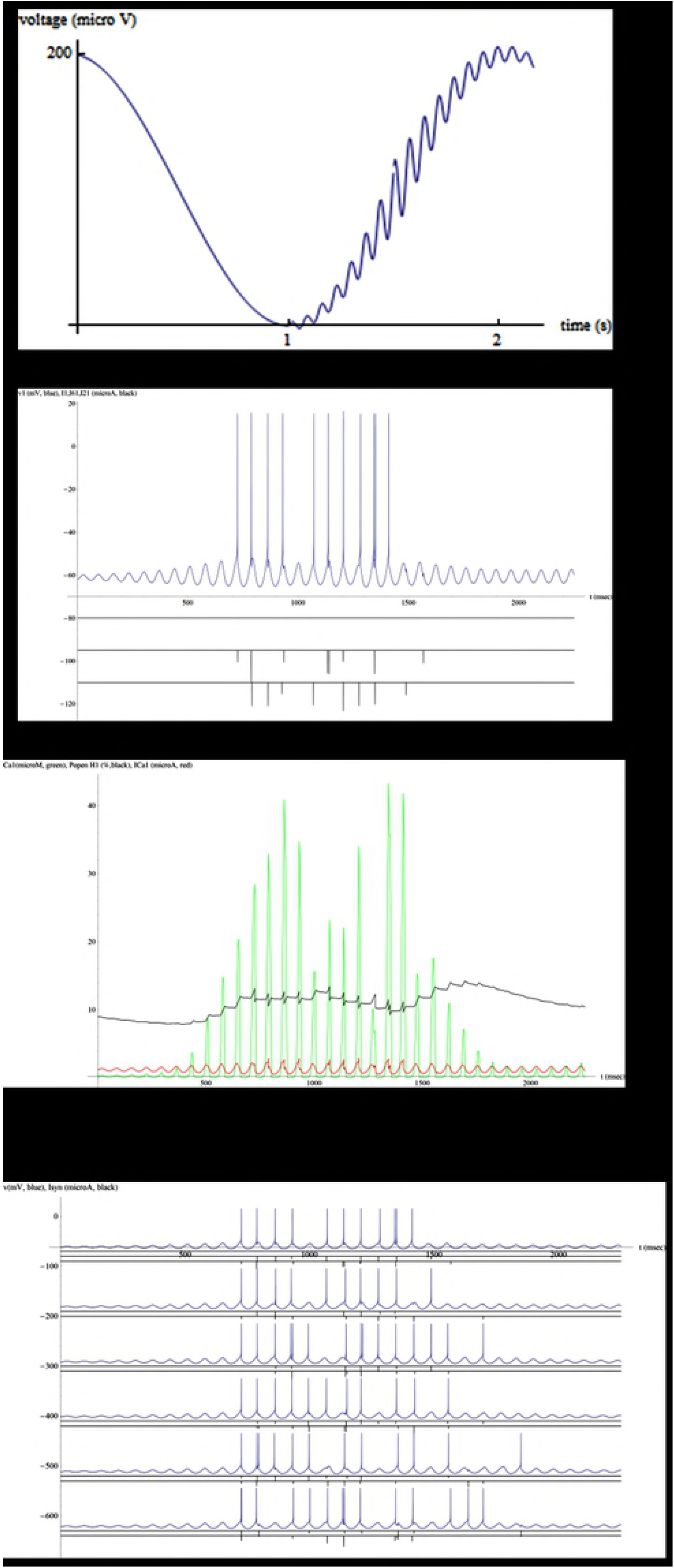
Self-excitation followed by spindle termination due to high calcium. A. Cartoon of a depth EEG recording of a local thalamic down state initiating a sleep spindle that exhibits waxing and waning of synchronized spikes between large numbers of relay neurons. B. Simulation of 6 relays poised at −61.3 mV slightly above the supercritical Hopf bifurcation (−61 mV). N1 shows a growing voltage oscillation and starts spiking at 720 ms, responding to EPSCs from N6 and N2. Calcium-clearance is low at 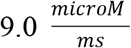and the calcium-sensitivity threshold of the H conductance low at 1.0 *microM*. C. Calcium (green) is able to raise the open probability of H conductance from 9 to 14% (black curve), depolarizing N1 and pulling it below the Hopf bifurcation. Increasing H conductance terminates the self-excited spindle. Kinks in H open probability identify N1 spikes, showing the competition between high calcium raising, and a voltage spike then lowering, its open probability. Small voltage oscillations only increase H open probability. Larger oscillations due to poising above the bifurcation make the spindle more sensitive to termination by calcium. D. Simulated spindle lasted 1.1 s with the relays exchanging 71 spikes. Average spike synchronization index is high (0.45). N3 and N4 spontaneously made action potentials. Presumably such a spiking focus would be able to entrain other relays poised near Hopf bifurcations having similar CFs.

Spindling relay neurons synchronize their spike bursts. In Fig. 4 we employ an event synchronization index [15] to quantify the amount of synch between them. We vary the delay between a spike in one relay neuron and its EPSC in a neighboring relay. Spikes in the group of relays are considered to be synchronized if they fall within 3 ms of each other. This is in the context of a subthreshold voltage oscillation with a 15 Hz CF and a 67 ms period in the relay cells. Synch and spike output drop with delays in the neighborhood of 1-2 ms. Plates 1C and 4B contrast examples of the high and low synch cases, respectively. Synch index drops by an order of magnitude in 4B’s 2.5 ms delay case (from 0.5 to 0.05).

**Fig.4.**
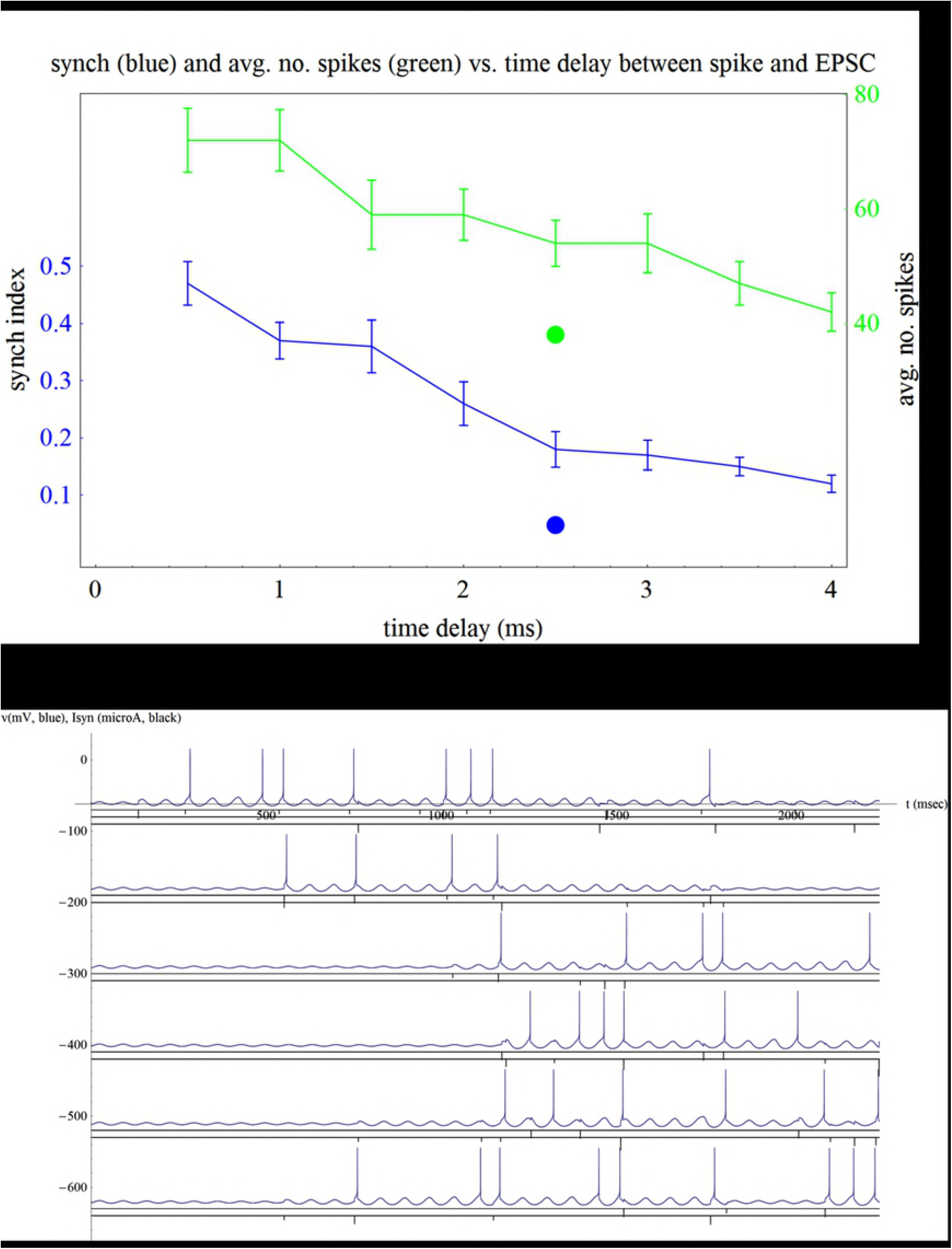
Spike synch as a function of the delay between a spike in a relay and its EPSC in a neighboring relay A. Simulations in other figures were run with short 0.5 ms delays between relay neurons. 1.0 ms delays had similar results, but with 20% lower synch. For delays > 1 ms synch and spike output drop significantly (average ±standard error). The period of the 15 Hz subthreshold oscillation is 67 ms and spikes were judged synchronized if they fell within 3 ms of each other (on the plateau of the voltage oscillation). B. Particularly low synch example caused by a 2.5 ms delay (dots in part A; same forcing, parameters as 0.5 ms delay high synch 1C).

The central claim of this study is that there are many advantages for poising relay neurons on supercritical Hopf bifurcations for the purposes of making and propagating well synchronized sleep spindles. Fig. 5 shows four of the main advantages: high small-signal gain, high degrees of frequency selectivity and noise rejection, and ease of switch-ability for turning receptivity on or off to a particular spindle. Plate A’s very high gain, calcium-insensitive case (purple) agrees well to the theoretical cube root shape for the forced response of a Hopf oscillator driven at its CF [11, 12]. As the sensitivity to calcium-induced depolarization of the neurons is increased (blue, green), spike gain predictably drops. It’s interesting that forcing of N1amounts to driving a nested sequence of Hopf oscillators, rather than a single one, as is done in hair cells in hearing. When a small number of current impulses force N1 at its natural frequency these lock at the optimal 90 degree phase lead with respect to the cell’s voltage oscillation. This is typical behavior for an electrical Hopf oscillator (mainly a capacitive load) when it’s poised at the bifurcation and responding with a high small signal gain [11, 12]. The other relay neurons then are forced by N1 and each other with much smaller < 10 degree phase leads and lags with respect to their limit cycle voltage oscillations. While this sacrifices gain, it preserves synch. However, when N1 is depolarized by *I_app_* bias current from −61 mV to below the Hopf bifurcation at its −60 mV high resting potential (5A, red curve), excitability goes well down. Spike output drops by an order of magnitude in the important weak forcing regime (compare to blue curve, same parameters, but with N1at −61 mV). So, by biasing the operating point of the input oscillator by ~1 mV the group of relay cells can be made either sensitive or insensitive to recruitment by a particular spindle. Also, output for aperiodic forcing (brown) is decreased by a factor of ~4 (compare to blue curve; same parameters, forcing). This shows the intrinsic noise rejection conferred by a Hopf oscillator.

**Fig.5.**
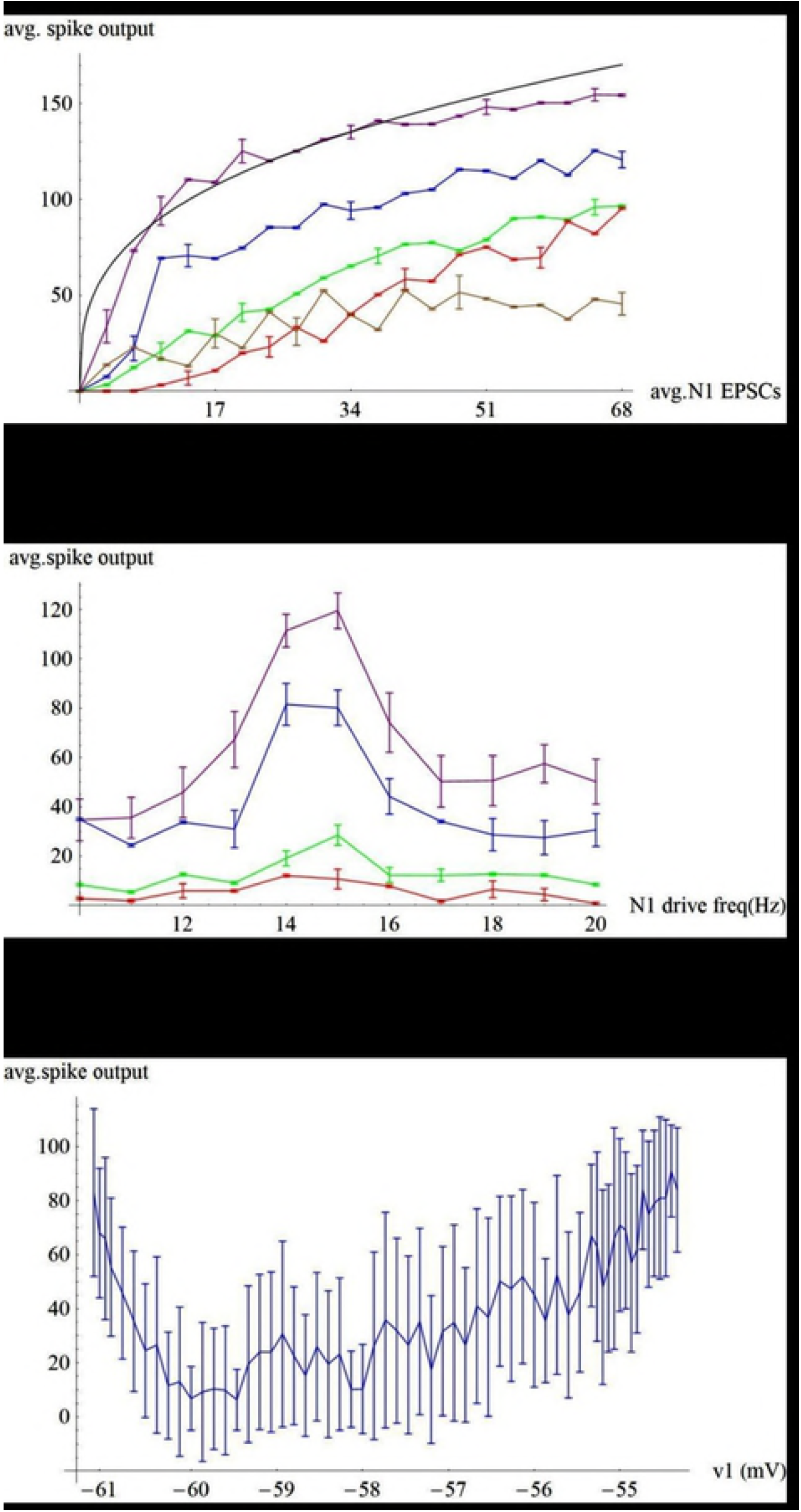
Advantages for poising relay neurons on supercritcial Hopfbifurcations. A. Hopf small signal amplification: total spike output as a function of the average number of current impulses driving N1 (average ± standard error). Poisson average vesicle release of 0.5 by an input spike averages 17 EPSCs forcing N1 during a 2.25 sec simulation (average release 1.0 results in 34, etc.). 17 impulses leads to 109 spikes on the ring of neurons in the high gain case (purple), which matches well to the theoretical cube root shape for the forced response of a Hopf oscillator driven at its CF (black). (*CaClear, CaSensH*, color; condition) = (10.0, 5.0, purple; max spike gain), (10.0, 2.0, blue; high gain), (10.0, 1.0, green; low gain), (10.0, 2.0, red; N1 poised at −60 mV, below the bifurcation), (10.0, 2.0, brown; aperiodic forcing of N1). B. Hopf frequency selectivity: spike output as a function of the N1 forcing frequency (same color scheme as A but with N1 forced by only 17 impulses). C. Switch-ability by altering a control parameter like N1 membrane potential. Spike output from periodic forcing at the CF by an average of 17 impulses driving N1 with it poised at various membrane potentials between the bifurcations (supercritical Hopf at −61 mV and saddle node on invariant circle (SNIC) at −55 mV). Spike output is very low when N1 is poised at its high resting potential (−60 mV). Spiking increases with depolarization from −59.5 to −55 mV. At these higher membrane potentials the low-threshold calcium conductance is completely off, but there is increasing activation of high-threshold sodium conductance, which puts the relay into the rate-coding region of its SNIC bifurcation. Spike output is noisy in this region so it was shown ± 1 standard deviation (instead of ± standard error for 20 runs).

5B shows Hopf small-signal frequency selectivity. The oscillators have a ~3 Hz full width half max (FWHM) band width when N1 is weakly forced in the 10-20 Hz range (purple, blue, green curves). But when N1 is depolarized by *I_app_* to its high resting state at −60 mV, band pass filtering is lost (red curve).

In 5C, N1 is weakly forced at its CF while being poised at various membrane potentials in between the bifurcations (supercritical Hopf at −61 mV and saddle node on invariant circle (SNIC) at −55 mV). There is a dead zone near the high resting state (−60 mV) where spike output bottoms. The high resting state sits above the activation range of the low-threshold calcium conductance and below the activation range of the high-threshold sodium conductance. In the voltage range between the bifurcations the input resistance is approximately 6 k Ohms so *I_app_* bias current of 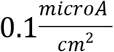displaces membrane potential by about 0.6 mV. Membrane potentials above the −60 mV high resting state move the cell closer to the SNIC, a bifurcation that is typically used by rate-coding sensory neurons [12]. Here the voltage range is used by thalamic relay cells acting in relay mode and being driven by subcortical sensory input (retina, cochlea, whiskers, etc.) to pass on sensory spikes in a mainly 1-1 manner [1]. In this rate-coding voltage range spike output in response to weak noisy periodic input is very noisy. In order to show the noise it was plotted ± 1 standard deviation, rather than the standard error for 20 runs.

To illustrate the nature of spike gain for very weak forcing, in Fig. 6 we show 4 short runs made under different conditions. For very weak on-frequency periodic input the current impulses (EPSCs) lock at a 90 degree phase lead with respect to N1’s voltage oscillation, increasing the size of its limit cycle until spikes result (A). But when N1 is depolarized close to its high resting potential at −60 mV, its rapidly decaying membrane potential oscillation is clear (B). This lowered excitability makes it hard to generate spikes. The reason for narrow Hopf frequency selectivity is clarified in plate C. Spike output drops sharply for input 2 Hz off CF because the current impulses are not forcing the voltage oscillation at the best phase. For impulsive currents the cell is mainly a capacitive load that is optimally forced at a 90 degree phase lead. For the same reason, weak aperiodic forcing is ineffective in generating spikes (D).

**Fig.6.**
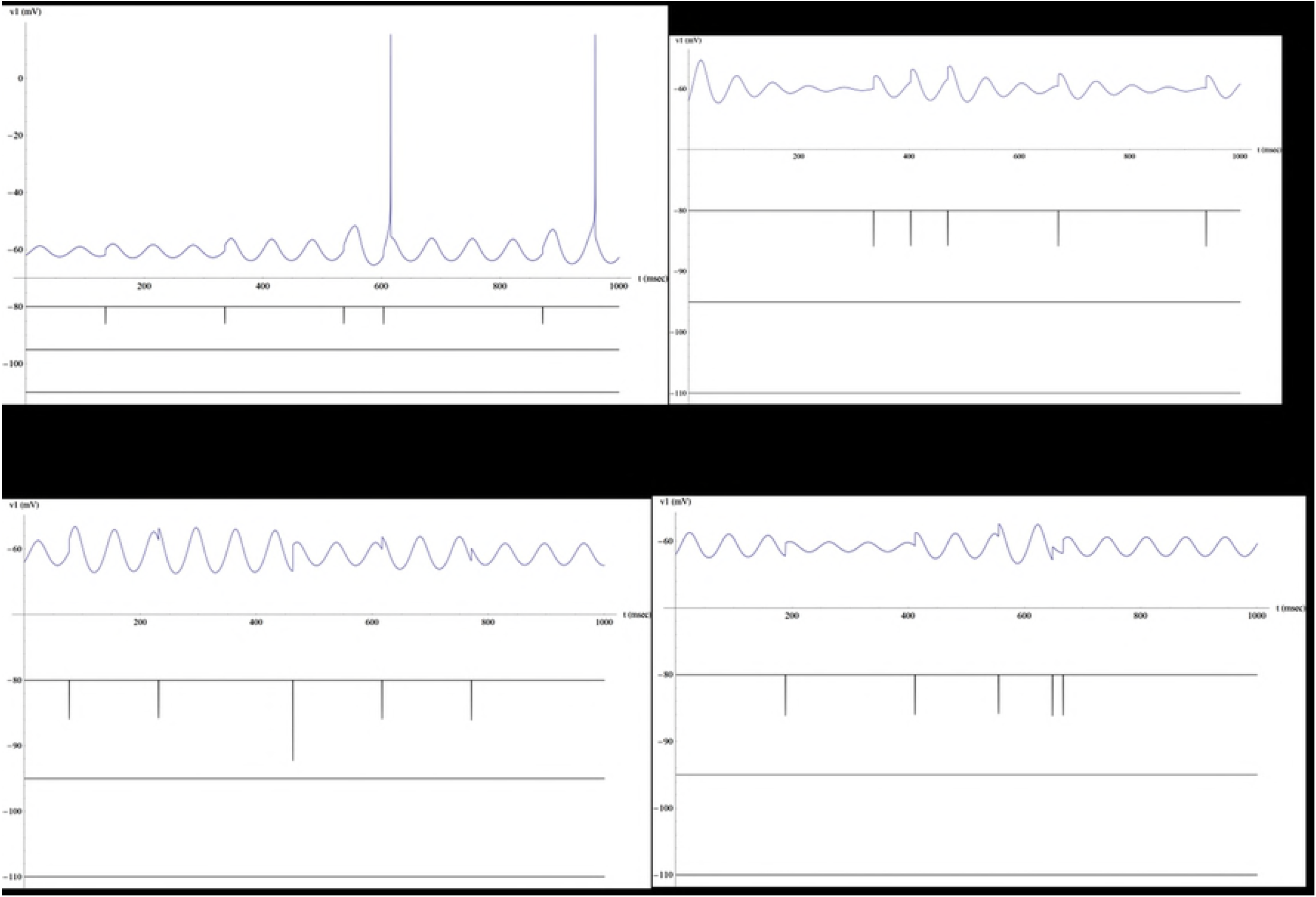
Short examples show spike gain for very weak external forcing of N1. A. Driving N1 at the CF (15 Hz) with it poised on the Hopf bifurcation at −61 mV. EPSCs lock at a 90 degree phase lead with respect to N1’s voltage response, increasing the size of its limit cycle oscillation until spikes result. B. Spikes are rare when forcing N1 at the CF, with it poised below the bifurcation in the dead zone at −60 mV. Note the rapidly decaying membrane potential oscillation. C. Spikes are rare when forcing N1 off CF at 13 Hz. D. Spikes rare for aperiodic forcing. 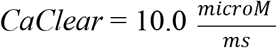, *CaSensH* = 2.0 *microM*.

It would be useful to have a visual low dimensional phase space representation for a relay neuron. The model neurons here have 6 dimensions: voltage (*v*), delayed-rectifier potassium channel activation gate (*n*), low-threshold calcium channel inactivation gate (*h*), calcium concentration (*Ca*), and the slower and faster calcium-, and voltage-, dependent H channel activation gates *H_s_* and *H_f_*. Very fast activation variables on the inactivating sodium conductance (*m*), persistent sodium conductance (*mp*) and low-threshold calcium conductance (*s*) are treated as instantaneous functions of voltage. The inactivation gate on the sodium conductance has a time course linearly related to the potassium conductance activation variable (.85-*n*) [9]. In this way, ten dynamical variables can be cut to six. Fig.7 A shows 11 EPSCs forcing N1 at the CF. Part B is an incomplete representation of the N1 phase space. Its voltage trajectory (*v1, h1, n1* taken from 520-750 ms in part A) shows 3 subthreshold oscillations, 2 EPSPs and one spike. Superposed in part C are 2 more incomplete representations (black; *v1, Ca1, H1*) and (orange; *v1, ICa1, H1*). In the subthreshold region, calcium current and concentration dovetail nicely. That being said, it’s hard to see how a relay neuron can be accurately represented by a low dimensional phase space representation.

**Fig.7.**
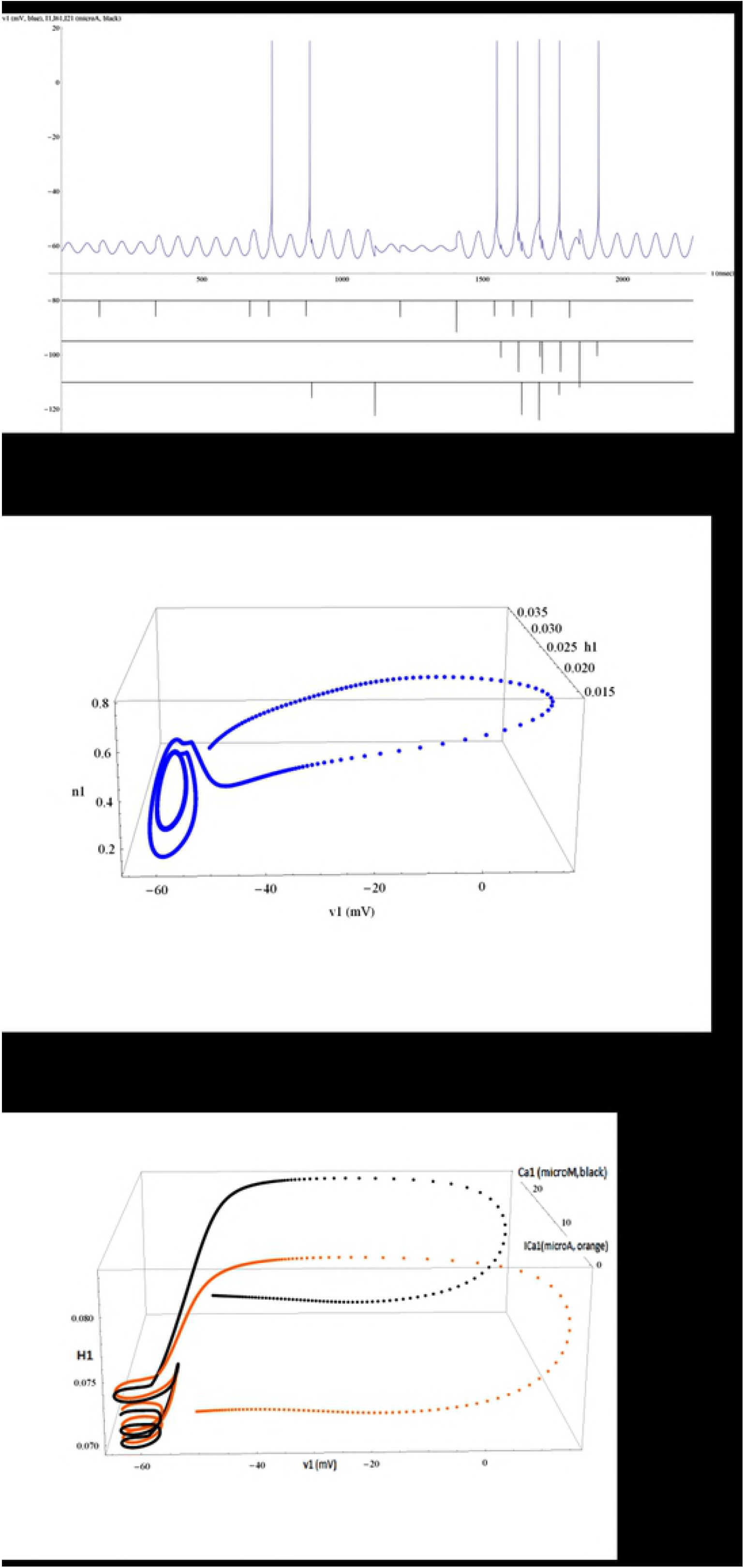
Reduced models of a thalamic relay neuron. A. N1 membrane potential (blue) and synaptic currents (black), external, N6 → N1 and N2 → N1 in descending order. B. 3D partial trajectory-style representation of N1: membrane potential *v1*, open probabilities of the potassium conductance activation gate *n1* and calcium conductance inactivation gate *h1*. Trace taken from 520-750 ms in part A shows 3 subthreshold oscillations, 2 EPSPs and a single spike. C. Two more trajectory-style representations from the same time segment. *v1*, calcium concentration *Ca1*, and H conductance open probability *H1* (black) and *v1*, calcium current *ICa1*, and *H1* (orange). 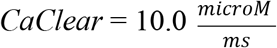, *CaSensH* = 1.0 *microM*.

## DISCUSSION

It’s interesting to look at the relative sizes and durations of the various currents in the relay neuron in order to see how these correlate with their particular functions. In this simple view each current is rated with an average strength and time interval: Linear leak current *I_L_* is about 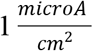 and is mainly outwards, continuous. Inward low threshold calcium current *I_T_* is about 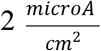 and persists for about 20 ms, during the upstroke of a subthreshold voltage oscillation. Inward *I_H_* is a continuous but very small bias current, about 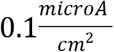. Calcium current *I_T_* and linear leak *I_L_* effectively run the subthreshold voltage oscillator (period ~ 67 ms), while *I_H_* is able to bias the size of its limit-cycle oscillation. Inward sodium currents *I_Na_* and *I*_*Na*(*P*)_ combine for a very large 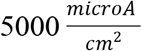, but only last for the upstroke of the action potential (about 100 micro sec). Outward delayed-rectifier potassium current *I_K_* goes about 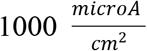 for 500 micro sec during the down stroke of the action potential. *I_S_* is an inward synaptic current of about 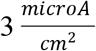 that lasts for 500 micro sec, forces a mainly capacitive load, and is able to make a 1-2 mV EPSP. *I_app_* stands in for various slower bias currents, and altering it by 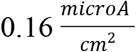 forces mainly a resistive load, and displaces the operating point by about 1 mV in the voltage range of interest (−60 to −61.2 mV; −60 mV = center of the range for the high voltage rest state; - 61 mV = supercritical Hopf bifurcation; −61.3 mV = fold-circle bursting bifurcation).

There are a number of advantages for spindling relay neurons to be poised on, or very near to, supercritical Hopf bifurcations: 1. Noise rejection, where weak aperiodic input results in low spike output. 2. Small signal amplification, combined with frequency selectivity, where forcing 2 Hz off CF halves spike output. 3. Tunable gain, where 3 parameters, the calcium clearance rate, the calcium-sensitivity threshold of the H current, and the membrane potential, effectively set spike gain. 4. Sleep spindles can be initiated by bias currents that poise a particular relay neuron’s membrane potential above the Hopf bifurcation. It will start to spontaneously spike at its natural frequency (CF) at the fold-circle bursting bifurcation. Spindles can then be propagated from this focus to other relays poised at Hopf bifurcations with a similar CF. 5. Switch-ability: susceptibility to a propagating sleep spindle can be turned off by depolarizing a relay to near its high resting potential of −60 mV. This decreases its small signal gain by an order of magnitude with respect to poising on the Hopf bifurcation at −61 mV. In this way a subnetwork of relay neurons can be switched into or out of participating in a particular sleep spindle. 6. Spindle termination can occur by calcium feedback activating H current that depolarizes the relay below the Hopf bifurcation.

Another advantage for poising on supercritical Hopf bifurcations is the idea of self-tuning, i.e. a self-tuned critical oscillation (STCO) [18, 19]. For example, Hopf amplifiers in hearing max their small-signal gain by seeking the bifurcation for their operating point. This has to do with the intrinsic nonlinearity in the sigmoidal Boltzmann channel activation curve. In the hearing case it involves the transduction channel [19]. In the relay neuron case it involves the *h* gate of the calcium conductance. Both turn off with depolarization, and below half open their slope conductance decreases with depolarization. The key point is that a small sinusoidal oscillation in membrane potential maps to an asymmetrical current oscillation which cycle-by-cycle generates a small negative feedback (net + current) that raises the membrane potential back to the point where the oscillation started, i.e. back to the bifurcation. Here, the *h* gate of the low-threshold calcium conductance has a low open probability and is poised in the lower knee region of its Boltzmann activation curve where the slope conductance changes most rapidly. So the net effect of a small voltage oscillation is to turn off the *h* gate on the calcium conductance. The upshot is that bias currents need only place the relay in the vicinity of the bifurcation, since it will then be attracted to it.

A spindle visible on an EEG lasts ½ to 2 sec. The early part of the spindle growth phase, involving few neurons, will be undetectable by EEG. Also, likely there are many smaller spindles which will never be detected by EEG. Here we model a 15 Hz spindle that lasts 10 to 30 cycles. In its decay phase calcium depolarizes the relays. Also, doublet, triplet bursting leads to phase-breaking. Both of these effects act to turn off the spindle. Sleep spindles are by necessity a kind of tickling of the dragon’s tail in that loss of their regulation would obviously be one way to get epilepsy. Therefore they must be focally initiated, strictly regulated, and terminated by strong negative feedbacks.

The “recall-store” memory consolidation hypothesis [2] envisions a 3 step process, with short term awake state memories alternately being spot-welded by spindles, then subsequently arc-welded in by delta and slow wave during NREM sleep. We argue that it is advantageous for the spindling relay neurons to be poised on supercritical Hopfbifurcations.

## REFERENCES

[1] Sherman MS (2005) thalamic relays and cortical functioning. Progress in Brain Research 149:107–124.

[2] Dextexhe A, Sejnowski TJ (2003) interactions between membrane conductances underlying thalamocortical slow wave oscillations. Physiol Rev 83:1401–1453.

[3] Andrillon T, Nir Y, Staba RJ, Ferrarelli F, Cirelli C, Tononi G, Fried I (2011) sleep spindles in humans: insights from intracranial EEG and unit recordings. J Neurosci 31(49): 17821–17834. DOI:10.1523/JNEUROSCI.2604-11.2011.

[4] Mak-McCully RA, Rolland M, Sargsyan A, Gonzalez C, Magnin M, Chauvel P, Rey M, Bastuji H, Halgren E (2017) coordination of cortical and thalamic activity during non-REM sleep in humans. Nat. Commun. 8:15499. doi: 10.1038/ncomms15499

[5] Tononi G, Massimini M, Riedner BA (2006) sleepy dialogues between cortex and hippocampus: who talks to whom. DOI:10.1016/j.neuron.2006.11.014

[6] Bartram J, Kahn MC, Tuohy S, Paulsen O, Wilson T, Mann EO (2017) cortical up states induce the selective3 weakening of subthreshold synaptic inputs. Nat Com 8:665. DOI:10.1038/s41467-017-00748-5

[7] Dextexhe A, Babloyantz A, Sejnowski TJ (1993) ionic mechanisms for intrinsic slow oscillations in thalamic relay neurons. Biophysical J 65: 1538–1552.

[8] Dextexhe A, McCormick DA, Sejnowski TJ (1993) model for 8-10 Hz spindling in interconnected thalamic relay and reticularis neurons. Biophysical J 65:2473–2477.

[9] Wang X-J (1994) multiple dynamical modes of thalamic relay neurons: rhythmic bursting and intermittent phase-locking. Neuroscience 59 (1): 21–31.

[10] Luthi A, McCormick DA (1998) H-current: properties of a neuronal and network pacemaker. Neuron 21: 9–12.

[11] Strogatz SH (1994) nonlinear dynamics and chaos: with applications to physics, biology, chemistry, and engineering. Perseus Books Publishing, LLC, Westview.

[12] Izhikevich EM (2000) neural excitability, spiking and bursting. International Journal of Bifurcation and Chaos 10(6): 1171–1266.

[13] Hobbie RK, Roth BJ (2007) intermediate physics for medicine and biology 4^th^ ed. Springer Science.

[14] Ashmore J (2010) the afferent synapse. In: The Oxford Handbook of Auditory Science: The Ear. Oxford University press. pp. 259–282.

[15] Quian Quiroga R, Kreuz T, Grassberger P (2002) event synchronization: a simple and fast method to measure synchronicity and time delay patterns. Phys Rev E 66(4) 1904–1913. DOI:191310.1103/PhysRevE.66.041904

[16] Allen C, Stevens CF (1994) an evaluation of causes for unreliability of synaptic transmission. PNAS 91:10380–10383

[17] Hackney CM, Furness DN (2010) hair bundle structure and mechanotransduction. In: The Oxford Handbook of Auditory Science: The Ear. Eds. Fuchs PA, Moore DR. Oxford University press. pp. 231–257.

[18] Julicher F (2003) active amplification by spontaneous hair bundle oscillations. In: Unsolved Problems of Noise and Fluctuations. Ed. Bezrukov SM. AIP Conference Proceedings 665: 109–115.

[19] Ospeck M, Iwasa KH (2003) noise in the outer hair cell and gain control in the ear. In: Unsolved Problems of Noise and Fluctuations. Ed. Bezrukov SM. AIP Conference Proceedings 665: 116–124.

